# Genomic signature of shifts in selection and alkaline adaptation in highland fish

**DOI:** 10.1101/2020.12.23.424241

**Authors:** Chao Tong, Miao Li, Yongtao Tang, Kai Zhao

## Abstract

Understanding how organisms adapt to aquatic life at high altitude is fundamental in evolutionary biology. This objective has been addressed primarily related to hypoxia adaptation by recent comparative studies, whereas highland fish has also long suffered extreme alkaline environment, insight into the genomic basis of alkaline adaptation has rarely been provided. Here, we compared the genomes or transcriptomes of 15 fish species, including two alkaline tolerant highland fish species and their six alkaline intolerant relatives, three alkaline tolerant lowland fish species and four alkaline intolerant species. We found putatively consistent patterns of molecular evolution in alkaline tolerant species in a large number of shared orthologs within highland and lowland fish taxa. Remarkably, we identified consistent signatures of accelerated evolution and positive selection in a set of shared genes associated with ion transport, apoptosis, immune response and energy metabolisms in alkaline tolerant species within both highland and lowland fish taxa. This is one of the first comparative studies that began to elucidate the consistent genomic signature of alkaline adaptation shared by highland and lowland fish. This finding also highlights the adaptive molecular evolution changes that support fish adapting to extreme environments at high altitude.

**Significance Statement:** Little is known about how wild fish responds to extreme alkaline stress besides hypoxia at high altitude. Comparative genomics has begun to elucidate the genomic basis of alkaline adaptation in lowland fish, such as killifish, but insight from highland fish has lagged behind. The common role of adaptive molecular evolution during alkaline adaptation in highland and lowland fish has rarely been discussed. We address this question by comparing 15 fish omics data. We find numbers of shared orthologs exhibited consistent patterns of molecular evolution in alkaline tolerant species relative to intolerant species. We further identify remarkably consistent signatures of rapidly evolving and positive selection in a substantial shared core of genes in both highland and lowland alkaline tolerant species.

## Introduction

Environments shape the genetic landscape of the populations that inhabit them (Witt & Huerta-Sánchez 2019). The Tibetan Plateau had experienced continuous uplift during the India-Asia collision since approximately 45 million years ago, that triggered numerous environmental changes (Li & Fang 1999; Favre et al. 2015). As elevation above sea level increases, a decrease in barometric pressure results in fewer oxygen molecules in the air, which causes hypoxia. Besides, other harsh environments highland wildlife have encountered include the long-term low temperature, and intensified ultraviolet radiation (An et al. 2001). Large numbers of endemic Tibetan animals had developed unique morphological, physiological or genetic features to tolerate such harsh conditions (Wen 2014; Tong, Tian, et al. 2017; Tong, Fei, et al. 2017). Basically, understanding how organisms adapt to extreme environment is fundamental to address many evolutionary questions, but it remains a formidable task to fully uncover the mechanism of adaptive process (Scheinfeldt & Tishkoff 2010; Tong, Fei, et al. 2017; Tong, Tian, et al. 2017). Adaptation at molecular level can occur by adaptive mutation in key genes over prolonged evolutionary time scales (Orr 2005). Recent studies employing genome-wide approaches have identified candidate genes associated with hypoxia and long-term cold response in Tibetan terrestrial wildlife adaptation to high altitude (Qu et al. 2013; Wu et al. 2020). Nevertheless, the draft genomes of very few Tibetan aquatic wildlife are sequenced (Xiao et al. 2020; Liu et al. 2019), the genomic basis of highland adaptation in aquatic animals (e.g. fish) remains largely unknown.

The schizothoracine fishes (Teleostei: Cyprinidae), the predominant fish fauna in the Tibetan Plateau, had evolved specific phenotypic characteristics to adapt to extreme aquatic environments, including hypoxia and long-term low temperature (Wu 1992; Cao et al. 1981). Recent comparative studies have identified key genes showing signals of positive selection during adaptation to such harsh environments (Yang et al. 2014; Wang et al. 2015; Kang et al. 2017), such as Hypoxia-inducible factor (HIF) (Guan et al. 2014) and Erythropoietin (EPO) (Xu et al. 2016) associated with hypoxia response, ATPase Family AAA Domain Containing 2 (ATAD2) (Tong, Fei, et al. 2017) and cAMP-dependent protein kinase catalytic subunit alpha (PRKACA) that involved into energy metabolism (Tong, Fei, et al. 2017). The main focus of previous studies in schizothoracine fishes are still on hypoxia and cold response. Notably, an increasing number of lakes in the Tibetan Plateau have been existing or towards alkaline due to the global climate changes and human activities (Zheng 1997). Thus, the increasing alkalization of fresh water has been the potential challenge to schizothoracine fishes. Among the schizothoracine fishes, *Gymnocypris przewalskii przewalskii* and *Gymnocypris przewalskii kelukehuensis* are the only two species inhabited extremely alkaline environment (Wu 1992). Unlike other broadly distributed schizothoracine fishes, such as *Gymnocypris eckloni, Schizopygopsis pylzovi* and *Platypharodon extremus* that inhabit in the Yellow river basin (Cao et al. 1981; Wu 1992; Qi et al. 2012), *G. p przewalskii* only inhabits in saline and alkaline lake. As the largest salt lake in China, Lake Qinghai (FIG. 1) is highly saline (up to 13‰) and alkaline (up to pH 9.4) water environment, a typical salt lake with unusually high sodium, potassium and magnesium concentration (Zheng 1997; Zhu & Wu 1975). In addition, *G. p. kelukehuensis* only inhabits in a soda lake located at the Tsaidam Basin in the northeastern Tibetan Plateau. Lake Keluke (FIG. 1) is also a soda lake with low salinity of 0.79‰ and high pH value up to 9.3 (Zheng 1997). Both schizothoracine fish species had developed unique physiological or genetic features to tolerate such harsh living conditions (Tong, Fei, et al. 2017). Therefore, this provides an exceptional model to investigate the genetic mechanisms underlying alkaline adaptation, and may provide novel insights to fully understand the mechanism of highland adaptation in fish as complement.

**FIG. 1.**
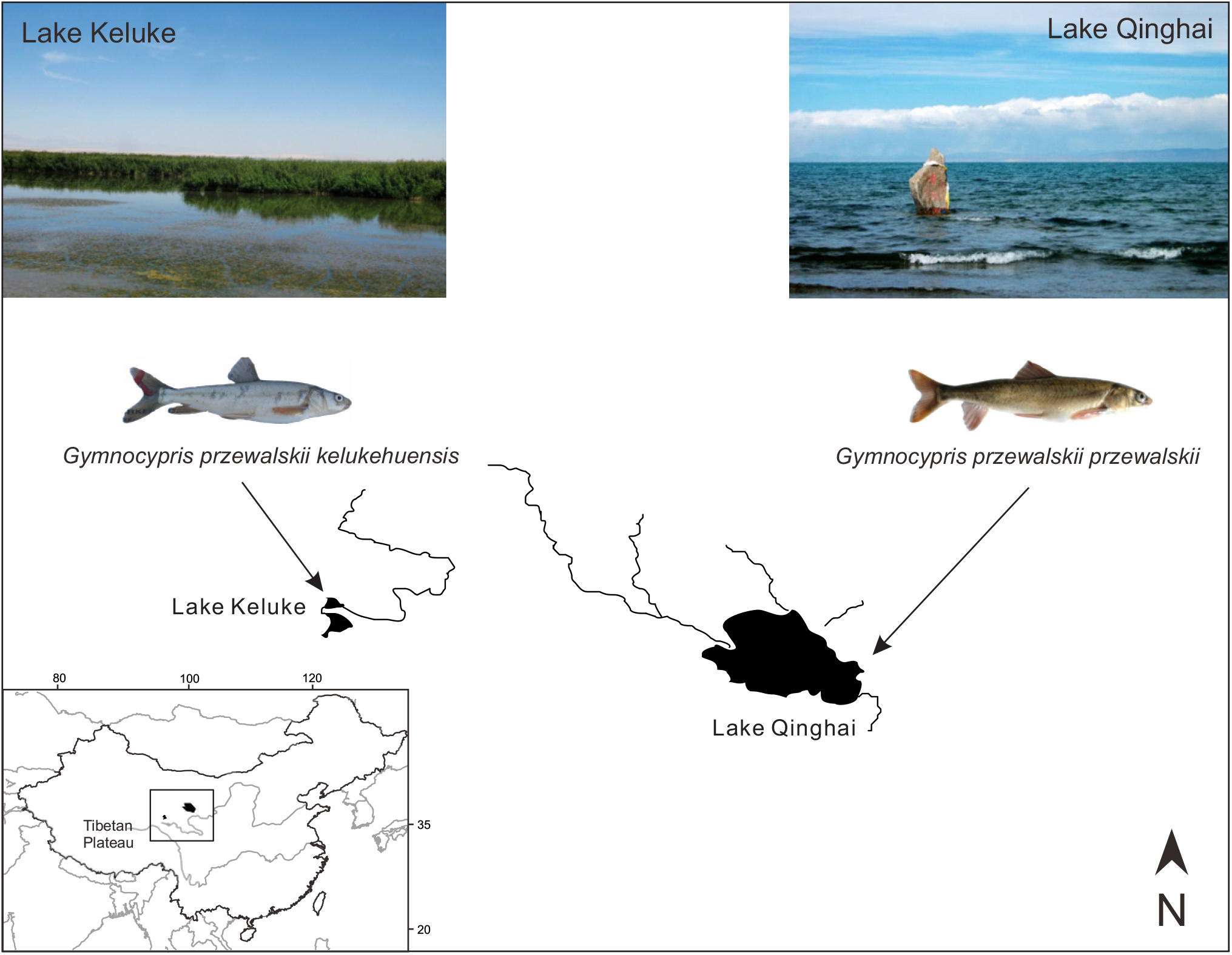
Overview of geographic distributions of two alkaline tolerant schizothoracine fish species and their habitats in the northeastern Tibetan Plateau. Map depicting the geographic distributions of *G. p. przewalskii* in Lake Qinghai, and *G. p. kelukehuensis* in Lake Keluke. Photos showing the representative specimens of *G. p. przewalskii* and *G. p. kelukehuensis*. Photo credit: Chao Tong and Kai Zhao.

Unlike highland alkaline tolerant fish, a huge amount of studies had explored the mechanisms of high saline and high alkaline tolerance in lowland fish species, such as killifish (e.g. *Fundulus heteroclitus*) (Wood et al. 2010;Brennan et al. 2018; Burnett et al. 2007), tilapia (e.g. *Oreochromis niloticus*) (Zhao et al. 2020; Wood et al. 1994), and salmonids (e.g. *Salmo salar*) (Levings 2016; Lien et al. 2016). These studies had provided insights in physiology of acid-base balance in fishes response to high salinity or high pH environments, and suggested key genes, such as ion transport associated genes under selection during the adaptation (Lien et al. 2016).

In this study, we generated and assembled the transcriptomes of two alkaline tolerant schizothoracine fish species, *G. p. przewalskii* and *G. p. kelukehuensis* inhabited in high pH environment in the northeastern Tibetan Plateau (FIG. 1). We performed a comparative genomics study together with recently sequenced schizothoracine fish transcriptomes and other lowland fish genomes (FIG. 2A & 2B), and sought to identify consistent genomic signature associated with alkaline adaptation in highland and lowland fishes. Specifically, we focused our comparisons on testing whether alkaline adaptation in highland and lowland alkaline tolerant fishes is associated with the following signatures of molecular evolution: (1) consistent patterns of molecular evolution in protein-coding genes across the phylogeny; (2) consistent shifts in evolutionary rates for specific genes; and (3) consistent signals of positive selection in particular genes.

**FIG. 2.**
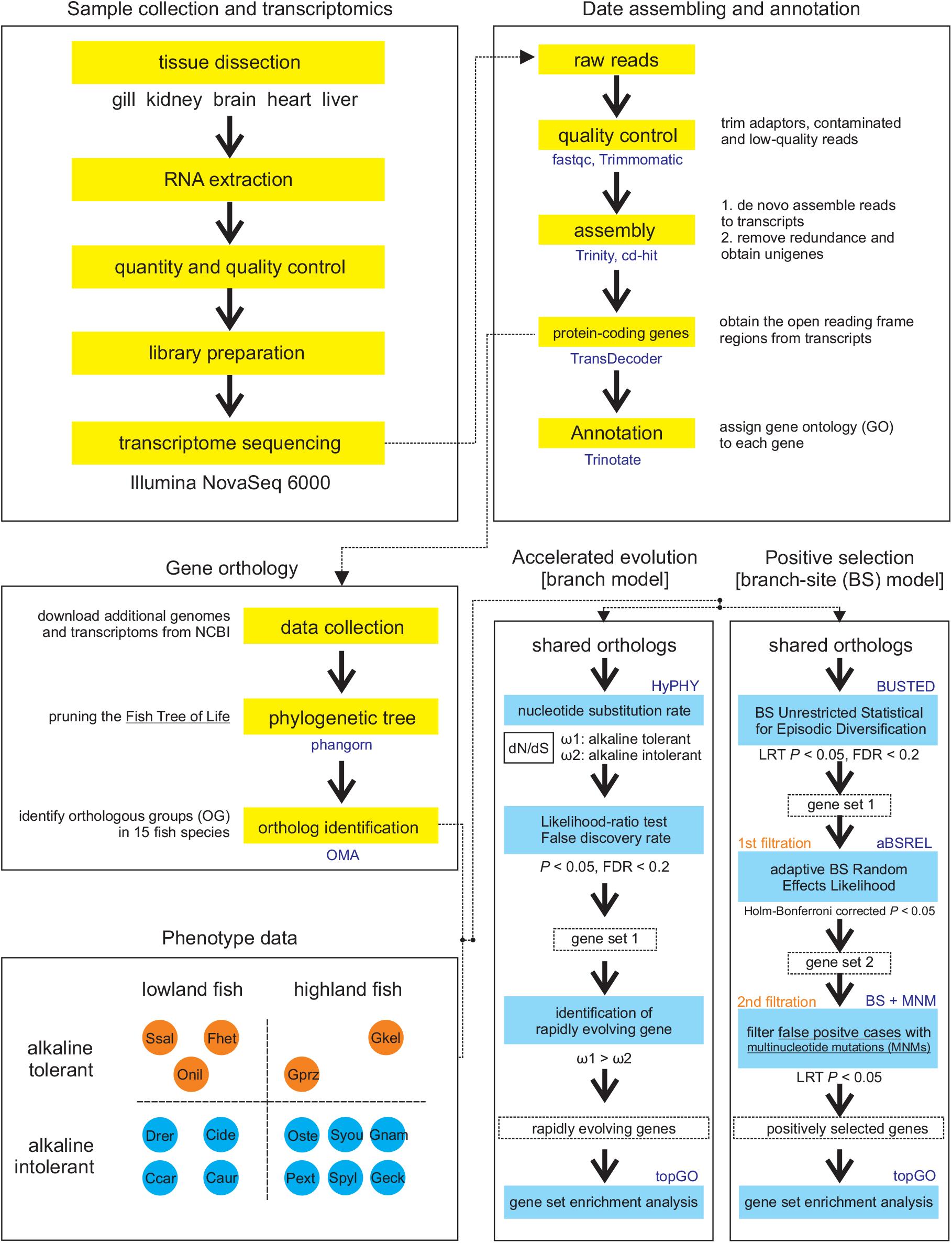
The flowchart represents the analysis pipeline: (1) sample collection and transcriptomics; (2) data assembling and annotation; (3) gene ortholog; (4) phenotype data and (5) molecular evolution analysis.

## Materials and Methods

### Sample collection

We collected eight adult *G. p. przewalskii* (FIG. 1) individuals (four males and four females, 172 ± 0.7 g) from Lake Qinghai and eight adult *G. p. kelukehuensis* (FIG. 1) individuals (four males and four females, 139 ± 0.3 g) from Lake Keluke using gill nets. All the fish samples were dissected after anesthesia with MS-222 (Solarbio, Beijing, China). All individuals were classified based on the gender and dissected after anesthesia with MS-222 (Solarbio, Beijing, China). Tissues from gill, kidney, brain, heart and liver from each individual were collected and immediately stored in liquid nitrogen at -80 °C. All the animal experiments were approved by the Animal Care and Use Committees of the Northwest Institute of Plateau Biology, Chinese Academy of Sciences (NWIPB-CY-010).

### Transcriptomics

Total RNA of each tissue sample was extracted using TRIzol reagent (Invitrogen, CA, USA) in accordance with manufacturer’s instructions, and detected for quality and quantity of RNAs with Nanodrop 1000 (NanoDrop Technologies, DE, USA) and Agilent Bioanalyzer 2100 (Agilent Technologies, CA, USA). Equal amount of RNA from eight individual of five tissue was pooled to construct transcriptome library as previously described (Tong, Tian, et al. 2017; Tong, Fei, et al. 2017), and sequenced with an Illumina NovaSeq 6000 yielding 150-bp paired-end reads (FIG.2).

Sequencing reads were checked for quality using FastQC (https://www.bioinformatics.babraham.ac.uk/projects/fastqc/). Sequencing adapters and reads with a quality score < 20 were trimmed with Trimmomatic (Bolger et al. 2014). We built a *de novo* transcriptome assembly based on clean reads using Trinity v2.6.5 (Grabherr et al. 2011) with default parameters. Next, we removed the redundant transcripts using CD-HIT (Fu et al. 2012) with the threshold of 0.90 and extracted the longest transcript as putative genes. We predicted the open reading frame of each putative genes using TransDecoder (https://github.com/TransDecoder/TransDecoder) (FIG.2).

### Additional data retrieval

We downloaded six alkaline intolerant schizothoracine fish transcriptomes (Zhou et al. 2020) including *Oxygymnocypris stewartii, Schizopygopsis younghusbandi, Gymnocypris namensis, Platypharodon extremus, Schizopygopsis pylzovi* and *Gymnocypris eckloni* from NCBI SRA database (https://www.ncbi.nlm.nih.gov/sra) (FIG. 3B, supplementary table S1), and performed assembly following above pipeline. In addition, we downloaded the genomes of four alkaline intolerant fish species of *Danio rerio, Ctenopharyngodon idellus, Cyprinus carpio*, and *Carassius auratus*, and three alkaline tolerant fish species of *Fundulus heteroclitus, Oreochromis niloticus* and *Salmo salar* (FIG. 3A, supplementary table S1).

**FIG. 3.**
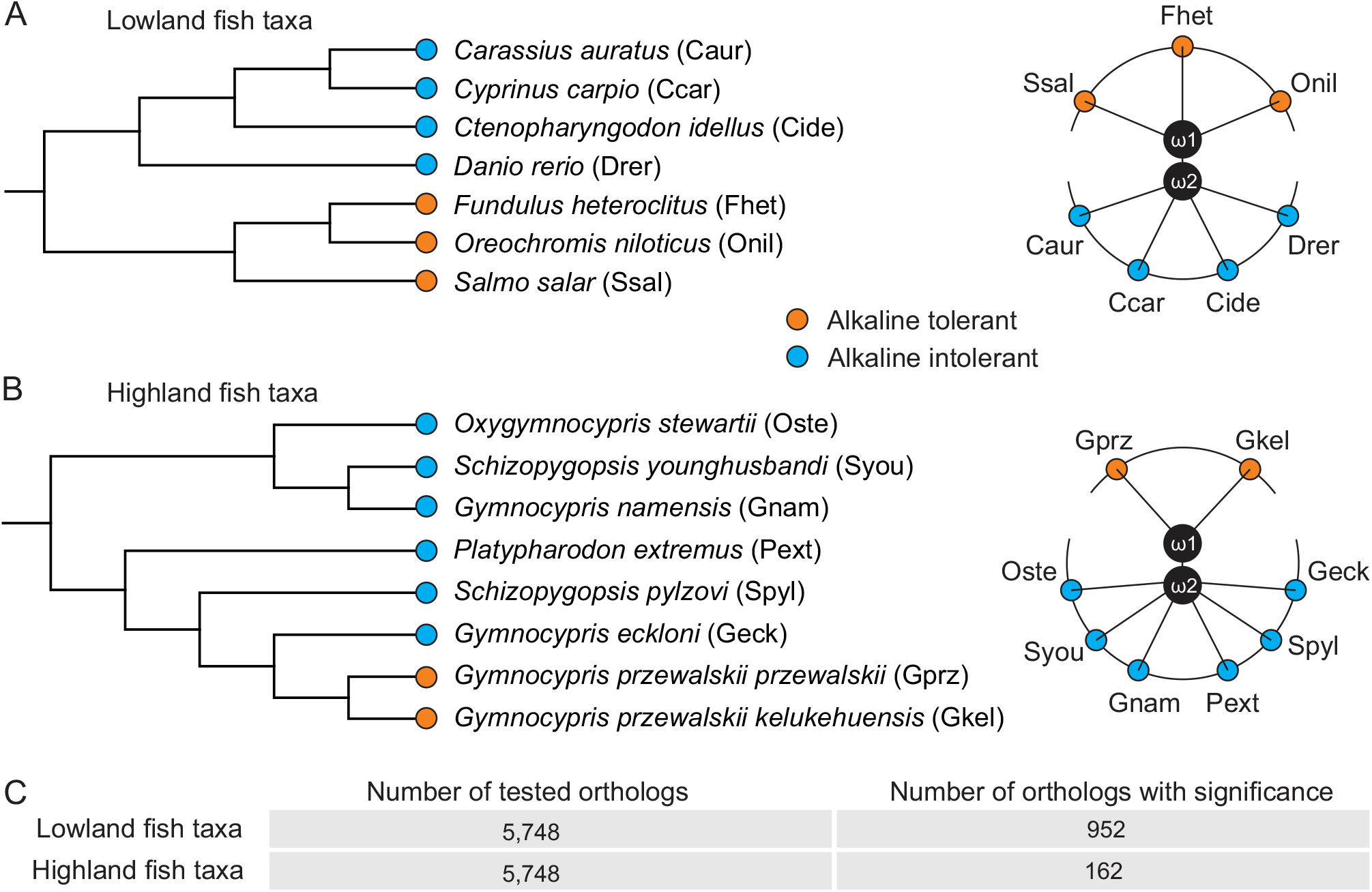
Consistent patterns of molecular evolution in alkaline tolerant fish species. (A-B) Species trees for lowland and highland fish taxa were pruned from the Fish Tree of Life (https://fishtreeoflife.org/). Alkaline tolerant taxa and alkaline intolerant taxa are depicted in orange and sky blue, respectively. Two schematic diagrams depicting the comparisons made between alkaline tolerant and alkaline intolerant fish species, ω1 representing the rate of molecular evolution of alkaline tolerant species and ω2 for the alkaline intolerant species (C) Number of tested orthologs (N = 5,748), and number of orthologs under consistent shifts in rates of molecular evolution in alkaline tolerant relative to alkaline intolerant species within highland fish taxa and lowland fish taxa (*P* < 0.05, likelihood ratio test; false discovery rate < 0.2).

### Species phylogeny and gene orthology

We obtained the phylogenetic tree of 15 fish species by pruning the Fish Tree of Life (https://fishtreeoflife.org/) using R package, phangorn (Schliep 2011). To obtain phylogeny-based orthology relationships between different fish taxa, we included all the predicted proteomes of seven lowland fish genomes, and translated nucleotide sequences of protein-coding genes from eight schizothoracine fish transcriptome assemblies into amino acid sequences, and pooled these datasets as input for an orthology inference tool, OMA (Altenhoff et al. 2018). In this way, we identified one-to-one, one-to-many, and many-to-many orthologs among these 15 fish species. For further comparison, we restricted our analysis to 1:1 orthologs, that is the geneset for which only one gene from each species representing the orthology. In addition, we extracted the shared orthologs among all fish taxa. At last, gene ontology (GO) terms were assigned to each ortholog using Trinotate (https://trinotate.github.io/) (FIG.2).

### Pattern of molecular evolution in shared orthologs

To determine whether highland fish and lowland fish showing consistent patterns of molecular evolution in the set of alkaline tolerant species branches across the phylogeny, we characterized the rates of non-synonymous to synonymous rate (dN/dS) in each shared ortholog. For this, we performed the protein sequence alignment using MUSCLE v3.8.31 (https://www.ebi.ac.uk/Tools/msa/muscle). We prepared the codon alignments of shared orthologs, that derived from protein alignments and the corresponding DNA sequences using PAL2NAL v.14 (Suyama et al. 2006). Then, we executed the filtration for shared ortholog alignments with length of at least 50 codons.

We took advantage of HyPHY pipeline (Kosakovsky Pond, Poon, et al. 2020) to test the hypotheses by comparing selective pressures (dN/dS) between *a prori* defined alkaline tolerant fish species branches (focal foreground branch) and alkaline intolerant species fish branches (background branch) in the specified fish phylogeny at ortholog-wide scale. Before that, a common approach to test the hypothesis is to perform separate analyses on subsets of sequences, and compare the parameter estimated in a *post hoc* fashion, that is one branch to the rest of branches in a phylogenetic tree like which we previously described (Tong, Tian, et al. 2017; Tong, Fei, et al. 2017; Tong et al. 2020). While, such approach is statistically suboptimal, and not always applicable (Kosakovsky Pond, Wisotsky, et al. 2020). In this way, we estimated two discrete categories of dN/dS for alkaline tolerant and alkaline intolerant species under the MG94 rate matrices combined with HKY85 model using HyPHY (Kosakovsky Pond, Poon, et al. 2020), and compared nested models with constrained relationships among them. We detected the shift of dN/dS ratios between alkaline tolerant and alkaline intolerant (alternative model, H1), relative to the null model that assuming all lowland fish taxa have the same dN/dS ratio (H0). We constructed the log-likelihood ratio score for each ortholog (ΔlnL) as: ΔlnL = 2(lnLH_1_-lnLH_0_) and employed the likelihood ratio test. Additionally, we applied a correction for multiple testing to a false discovery rate (FDR) < 0.20 using R package, q-value (http://github.com/jdstorey/qvalue). In this way, we repeated this analysis with shared ortholog datasets of highland taxa.

### Analysis of accelerated evolution

To determine if consistent shift in evolutionary rates (e.g. acceleration) in particular genes within alkaline tolerant species across the phylogeny, we defined rapidly evolving genes with significantly higher dN/dS ratio for alkaline tolerant species than alkaline intolerant species (*P* < 0.05, likelihood ratio test, FDR < 0.2). In this way, we identified the rapidly evolving genes in alkaline tolerant species within highland and lowland based on earlier estimated dataset including two discrete dN/dS ratios for each shared ortholog, respectively. In addition, these sets of rapidly evolving genes were examined for GO enrichment relative to the full set of all shared orthologs using R package, topGO (https://bioconductor.org/packages/release/bioc/html/topGO.html). We finally visualized all significantly enriched GO terms remaining after the REVIGO (http://revigo.irb.hr/) similarity filter.

### Analysis of positive selection

To further determine whether consistent signals of positive selection in a set of branches representing alkaline tolerant species across the phylogeny in specific genes, we used three complementary branch-site models to identify the positively selected genes (FIG.2). Specifically, we first used the Branch-Site Unrestricted Statistical Test for Episodic Diversification (BUSTED) model (Murrell et al. 2015) to test for positive selection signal in a gene at any site on focal branches. In this model, it defines a positively selected gene (LRT *P* < 0.05, FDR < 0.2) with at least one site under positive selection on at least one focal foreground branch, while does not specify the exact branch with positively selected site, may include false positive. Then, we used an adaptive Branch-Site Random Effects Likelihood (aBSREL) modelaBSREL (Smith et al. 2015) to test if positive selection has occurred on a proportion of branches (i.e. number of focal foreground branches) at specific genes (Holm-Bonferroni corrected *P* < 0.05). This allowed us to filter the false positive cases without positive selection signal on specific focal foreground branches (i.e. alkaline tolerant fish) out of the gene sets determined by BUSTED (FIG.2).We defined the positively selected genes that required the shift across all focal foreground branches. Since the branch-site models can cause false positive in case of multinucleotide mutations (MNMs, Venkat et al. 2018), we performed more conservative branch-site (BS) model test covering MNM situation (BS + MNM) (github.com/JoeThorntonLab/MNM_SelectionTests). In this model, an additional parameter δ represents the relative instantaneous rate of double mutations compared to that of single mutations. We ran null models and alternative models in BS + MNM and conducted LRTs to evaluate significance (LRT *P* < 0.05). In this way, we further filter false positive cases out of the previously positively selected gene sets that determined by BUSTED and aBSREL (FIG.2). In this way, we took advantage of these three models to finalize the positively selected genes (i.e. gene with sites under positive selection) in alkaline tolerant species within highland fish taxa and lowland fish taxa, respectively (FIG.2). We finally performed GO enrichment analysis using topGO and REVIGO as earlier described.

### Intersection of rapidly evolving and positively selected gene repertoire

To determine if there is both accelerated evolution and positive selection for specific genes, we did the overlapping between rapidly evolving genes (REGs) and positively selected genes (REGs) for highland and lowland alkaline tolerant fish species. In addition, we performed GO enrichment analysis with topGO and REVIGO for the overlapping genes.

## Results

Following *de novo* assembly and annotation, each of the eight schizothoracine fish transcriptome assemblies had an average of 39,921 transcripts representing the complete or partial protein-coding regions of genes (supplementary table S2). We further identified a total of 7,309 one-to-one orthologs shared by 15 fish species (i.e. range from 2 to 15 fish species), and 6,241 shared 1:1 orthologs including all 15 species (supplementary table S3).

### Consistent pattern of molecular evolution

We estimated two categories of dN/dS ratios for 5,748 shared 1:1 orthologs (after restricting to 1:1 orthologs with at least 50 codons) in lowland fish taxa and highland fish taxa, separately. We found consistent patterns of molecular evolution showing significant (LRT, *P* < 0.05,FDR < 0.2) acceleration (increased dN/dS) or deceleration (decreased dN/dS) in a set of terminal branches of alkaline tolerant species relative to the set of terminal branches of alkaline intolerant species in large numbers of shared orthologs within lowland fish taxa (n = 952) and highland fish taxa (n = 162) (FIG. 3C, supplementary table S4).

### Consistent signature of accelerated evolution

Building on earlier dataset of two discrete categories of dN/dS ratios separately representing branches of alkaline tolerant and alkaline intolerant species across the phylogeny, we focused on genes with significantly higher dN/dS (LRT, *P* < 0.05,FDR < 0.2) in alkaline tolerant species, that is rapidly evolving gene (REGs) repertoire in alkaline tolerant species within highland and lowland fish taxa. We identified 110 REGs in highland fishes and 470 REGs in lowland fishes (FIG. 4A). Out of 11 overlapping REGs, we found a set of ion transport and transmembrane functions associated genes, such as sodium-dependent phosphate transport protein 2A (SLC34A1) and sodium-dependent phosphate transporter 1-B (SLC20A1b) (FIG, 4B, supplementary table S5). Besides, we identified overlapping REGs related to energy metabolism process, such Ectonucleotide pyrophosphatase/phosphodiesterase family member 1 (ENPP1) (FIG. 4B). Given that different REGs identified in either highland or lowland alkaline tolerant fish species, we found a number of ion transport and ATP synthesis related genes in REGs repertoire of highland fish (FIG. 4B, supplementary table S5), such as solute carrier family 35 member F4 (SLC35F4), ATP-sensitive inward rectifier potassium channel 15 (KCNJ15), sodium-dependent serotonin transporter (SLC6A4), and NADP-dependent malic enzyme (ME1). Similarly, in lowland fish, we also identified genes like solute carrier family 45 member 3 (SLC45A3), Potassium Inwardly Rectifying Channel Subfamily J Member 5 (KCNJ5), and ATP synthase F0 complex subunit B1 (ATP5PB) showing rapidly evolving in alkaline tolerant species (FIG. 4B, supplementary table S5).

**FIG. 4.**
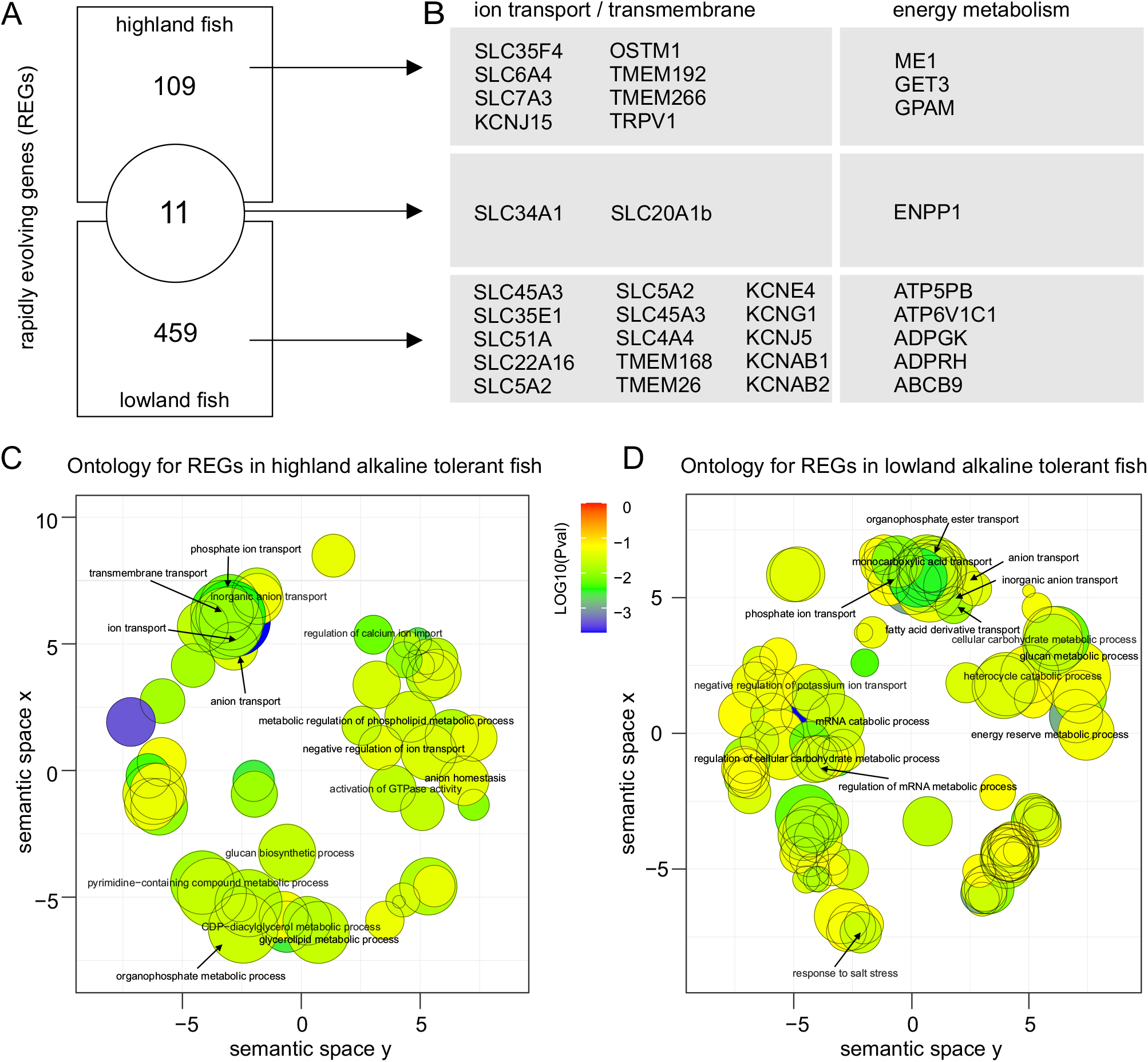
Consistent signature of accelerated evolution in alkaline tolerant species within highland and lowland fish taxa. (A) Venn diagram depicting numbers of rapidly evolving genes (REGs) in highland alkaline tolerant species, lowland alkaline tolerant species and both. (B) Highland fish specific REGs, lowland fish specific REGs and overlapping REGs mainly related to ion transport, transmembrane and energy metabolism functions. The table showing the representative REGs under each category. (C) REVIGO plot depicting the dominant enriched GO terms for REGs in highland alkaline tolerant fish. The scale of dark dot indicating the number of included enriched GO terms under a dominant GO term, the color scale representing the *P* value transformed by log10. (D) REVIGO plot depicting the dominant enriched GO terms for REGs in lowland alkaline tolerant fish.

Further, we did GO enrichment analysis for both REGs datasets, showing 112 significantly enriched GO terms (biological process) in highland fish (*P* < 0.05, fisher’s exact test) and 189 significantly enriched GO terms in lowland fish. After the filtration by semantic similarity of GO terms, we found a set of enriched dominant GO terms in highland fish were related to ion transport and transmembrane functions (FIG. 4C, supplementary table S6), such as ion transport (GO:0006811), regulation of calcium ion transport (GO:0051924), anion transport (GO:0006820). In addition, in lowland fish, we observed a set of enriched dominant GO terms associated with ion transport function (FIG. 4D, supplementary table S6), such as phosphate ion transport (GO:0006817) and response to salt stress (GO:0009651). Finally, we found 5 overlapping GO terms related to ion transport and metabolism, including glycogen biosynthetic process (GO:0005978), phosphate ion transport (GO:0006817), inorganic anion transport (GO:0015698), negative regulation of ion transport (GO:0043271), regulation of glycogen metabolic process (GO:0070873) and anion transport (GO:0006820) (supplementary table S6). Collectively, this finding suggested the consistent signature of accelerated evolution in alkaline tolerant species within highland taxa and lowland taxa.

### Consistent signature of positive selection

After two rounds of filtration with the use of BUSTED, aBSREL and BS+MNM models, we identified 162 positively selected genes in highland alkaline tolerant fish (FIG. 5A, supplementary table S7), and 156 PSGs in lowland alkaline tolerant fish species (FIG. 5A, supplementary table S7). Out of 7 overlapping PSGs, we found these genes were mainly related to apoptosis (cell death), ion transport and immune response, such as vitamin K-dependent protein C precursor (PROC), E3 ubiquitin-protein ligase SH3RF1 (SH3RF1), 14-3-3 protein beta/alpha-A (YWHABA) and cadherin-related family member 2 (Cdhr2). Besides, we found PSGs in highland alkaline tolerant species were also mainly involved in similar functional categories as overlapping PSGs (FIG. 5B, supplementary table S7), such as transmembrane protein 268 (TMEM268), transmembrane protein 266 (TMEM266), solute carrier family 35 member F4 (SLC35F4), solute carrier family 7, member 3 (SLC35F4) and solute carrier family 7, member 3 (SLC7A3) involved in ion transport or transmembrane functions, probable phospholipid-transporting ATPase VD (ATP10D) involved in energy metabolism, apoptosis-inducing factor 1 (AIFM1) involved in apoptosis, interleukin-2 receptor subunit beta (IL2RB) related to immune response. Similarly, in lowland alkaline tolerant fish species, these PSGs were involved in four main categories (FIG. 5B, supplementary table S7), such as solute carrier family 2 member 15b (SLC2A15b and potassium voltage-gated channel subfamily E member 4 (KCNE4) associated with ion transport function, phosphoinositide 3-kinase regulatory subunit 4 (PIK3R4) and lysosomal-associated membrane protein 1 (LAMP1) associated with apoptosis, CD22 antigen (CD22) and immunoglobulin-like domain containing receptor 1b precursor (ILDR1) involved in immune response, and plasma membrane calcium-transporting ATPase 3 (ATP2B3) associated with energy metabolism.

**FIG. 5.**
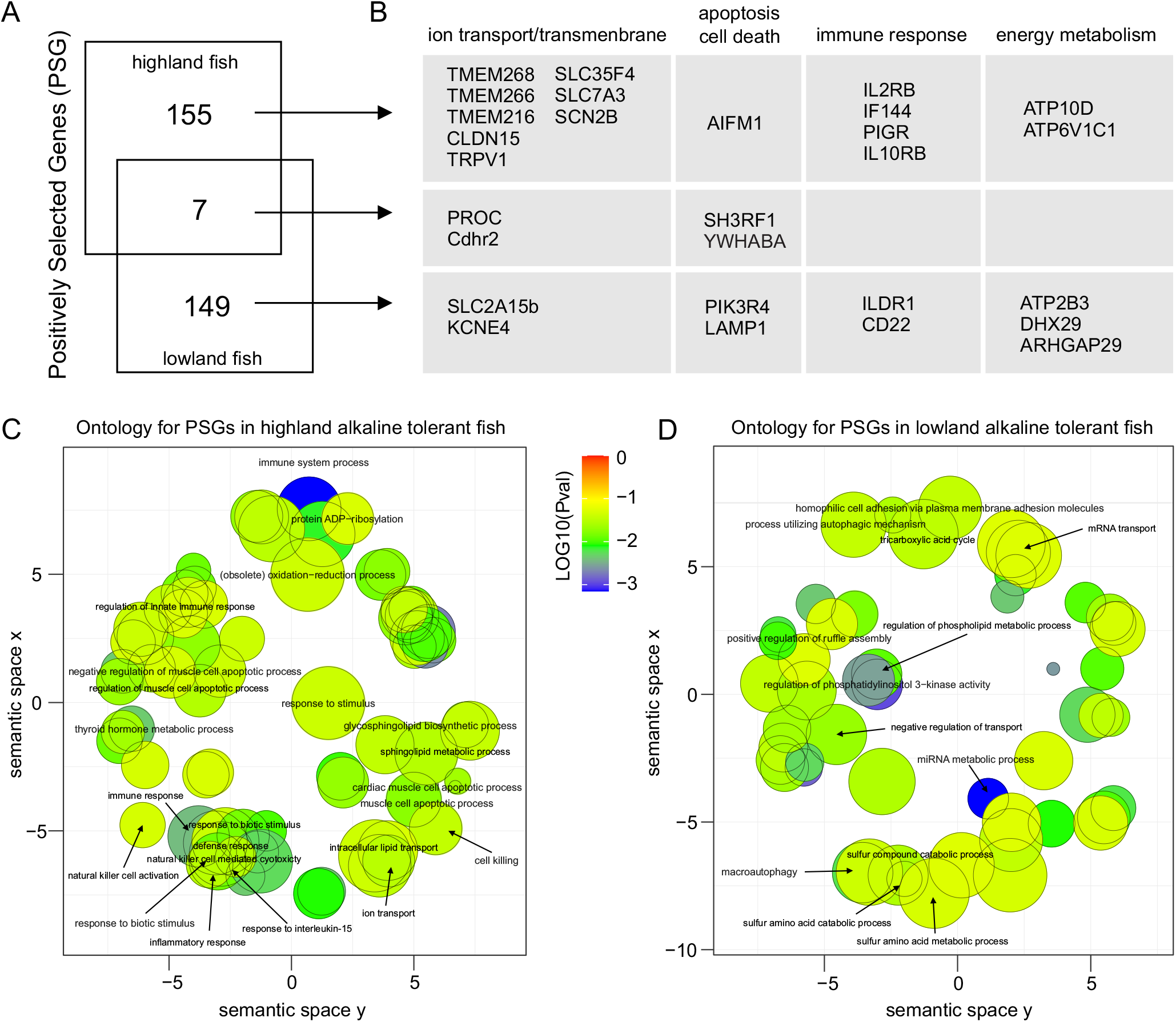
Consistent signature of positive selection in alkaline tolerant species within highland and lowland fish taxa. (A) Venn diagram depicting numbers of positively selected genes (PSGs) in highland alkaline tolerant species, lowland alkaline tolerant species and both. (B) Highland fish specific PSGs, lowland fish specific PSGs and overlapping PSGs mainly related to four categories, including ion transport/transmembrane, apoptosis/cell death, immune response and energy metabolism. The table showing the representative PSGs under each category. (C) REVIGO plot depicting the dominant enriched GO terms for PSGs in highland alkaline tolerant fish. The dark dot scale indicating the number of included GO terms under a dominant GO term, the color scale representing the *P* value transformed by log10. (D) REVIGO plot depicting the dominant enriched GO terms for PSGs in lowland alkaline tolerant fish.

Further, GO enrichment results showed that a number of significantly enriched GO terms (*P* < 0.05, fisher’s exact test) are related to immune response, apoptosis, protein metabolisms for PGSs in highland alkaline tolerant species, such as inflammatory response (GO:0006954), natural killer cell activation (GO:0030101), cell killing (GO:0001906), muscle cell apoptotic process (GO:0010656), regulation of innate immune response (GO:0045088), immune response (GO:0006955), ion transport (GO:0006811), sphingolipid metabolic process (GO:0006665) and thyroid hormone metabolic process (GO:0042403) (FIG. 5C, supplementary table S8). Similarly, in lowland alkaline tolerant species, the significantly enriched GO terms were mainly related to four categories as well, such as T cell mediated cytotoxicity (GO:0001913) related to immune response, negative regulation of transport (GO:0051051) and negative regulation of ion transport (GO:0043271) related to transport function, macroautophagy (GO:0016236) related to apoptosis (cell death), regulation of phospholipid metabolic process (GO:1903725) and sulfur amino acid metabolism (GO:0000096) (FIG. 5D, supplementary table S8). Collectively, this finding suggested the consistent signature of positive selection in alkaline tolerant species within highland taxa and lowland taxa.

### Genes with evidence for both accelerated evolution and positive selection

We sought to find the intersection of REGs and PSGs repertoires, we found 29 overlapping genes in highland alkaline tolerant fish (FIG.6A), mainly related to ion transport, apoptosis (cell death), and energy metabolism, such as SLC35F4, SLC7A3, SLC6A4, TMEM266, CLDN15 and ALDH16A1 and ATP6V1C1 (supplementary table S9). GO enrichment also suggested that overlapping genes were mainly enriched in three main functional categories, such ion transport (GO:0006811), anion transport (GO:0006820), regulation of calcium ion import (GO:0090279), alditol phosphate metabolic process (GO:0052646), oxoacid metabolic process (GO:0043436) and regulation of muscle cell apoptotic process (GO:0010660) (FIG.6B, supplementary table S10). Similarly, we observed 26 shared REG/PSG genes in lowland alkaline tolerant fish (FIG. 6C, supplementary table S9). The additional GO enrichment result showed that the overlapping genes mainly enriched into transport and metabolic process, such as negative regulation of ion transport (GO:0043271), negative regulation of potassium ion transmembrane transporter activity (GO:1901017) and GTP metabolic process (GO:0046039) (FIG. 6D, supplementary table S10). Thus, this finding emphasized the consistent signatures of accelerated evolution and positive selection in alkaline tolerant species.

**FIG. 6.**
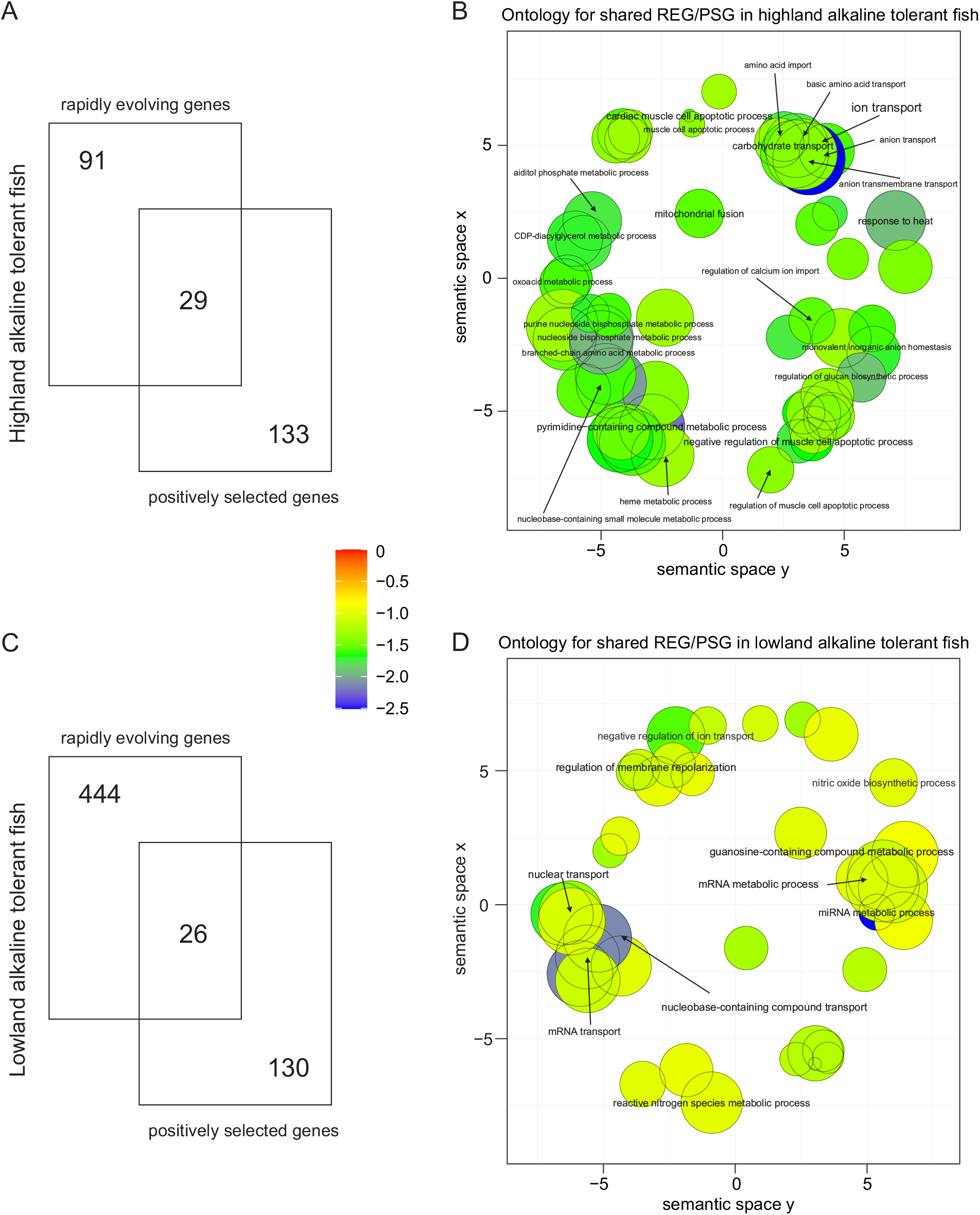
The intersection of rapidly evolving genes (REGs) and positively selected genes (PSGs) in alkaline tolerant species within highland and lowland fish taxa. (A) Venn diagram depicting numbers of REGs, PSGs and their overlapping genes in highland alkaline tolerant species. (B) REVIGO plot depicting the dominant enriched GO terms for overlapping genes in highland alkaline tolerant species. (C) Venn diagram depicting numbers of REGs, PSGs and their overlapping genes in lowland alkaline tolerant species. (D) REVIGO plot depicting the dominant enriched GO terms for overlapping genes in lowland alkaline tolerant species.

## Discussion

Our results support above three hypotheses that alkaline tolerant species shared the consistent patterns of molecular evolution in protein-coding genes (dN/dS) and consistent signatures of accelerated evolution (rapidly evolving gene) and positive selection (positively selected gene) in highland and lowland fish. Specifically, these signatures include: genes experienced consistent acceleration in evolutionary rates (increased dN/dS) in alkaline tolerant species, which are mainly involved in ion transport, transmembrane and energy metabolism functions; genes showing consistent signals of positive selection in alkaline tolerant species within highland and lowland fish taxa, these are mainly associated with ion transport/transmembrane, apoptosis (cell death), immune response and energy metabolism processes. Altogether, this study provides insights in understanding the common role of adaptive molecular evolution in fish adaptation to alkaline environment, as well as adaptation to extreme environment at high-altitude.

### Acid-base balance and osmoregulation

In freshwater fish, Na^+^ and Cl^−^ are actively taken up across the gill epithelium to counter the passive loss of osmolytes to the more dilute environment. After transition to saline or alkaline water, fish must increase its rate to balance the osmotic loss of water to the more solute-concentrated environment and actively excrete Na^+^ and Cl^−^ from the gill to maintain ionic and osmoregulation (Marshall 2005). Thus, alkaline tolerant species requires enhanced physiological abilities including acid-base balance and osmoregulation to respond the elevation in alkalinity or salinity of freshwater (Evans et al. 2005; Marshall 2005). Extremely alkaline environment may accelerate the evolution of genes associated with osmoregulation in these species survived in such harsh environment (Xu et al. 2017; Tong, Fei, et al. 2017). In this study, we identified a set of genes associated with ion transport and transmembrane that tended to evolve rapidly in alkaline tolerant species than their alkaline intolerant relatives (FIG. 4). This result echoed our previous finding in *G. przewalskii* compared with other teleost fishes (alkaline intolerance) (Tong, Fei, et al. 2017), also was in line with the rapidly evolving gene repertoire of a wild fish, Amur ide (*Leuciscus waleckii*) that survived in extremely alkaline environment (Xu et al. 2017). Interestingly, osmoregulation related genes including three solute carrier (SLC) genes and one transient receptor potential cation channel (TRPV) gene exhibited consistent signature of accelerated evolution in both highland and lowland alkaline tolerant fishes. *SLC* genes encode transmembrane transporters for inorganic ions, amino acids, neurotransmitters, sugars, purines and fatty acids, and other solute substrates (Dorwart et al. 2008). Recent evidences indicted that adaptive evolution of *SLC* genes contributed to fish adaptation to high salinity and high pH environment (Wang & Guo 2019; Tong, Fei, et al. 2017; Xu et al. 2017). Besides, we also identified numbers of ion transport and transmembrane genes under positive selection in alkaline tolerant fish species, such as potassium voltage-gated channel (*KCN*) genes. In killifish, an excellent model to study extreme environment adaptation, recent genome-wide studies also found *KCN* genes have been implicated in freshwater adaptation (transition from marine to freshwater environment) (Brennan et al. 2018).) Besides a set of same rapidly evolving and positively selected genes identified in both highland and lowland alkaline tolerant fish species, we found a large number of different genes under selection but involved in similar functions, such as anion transport (GO:0006820) and phosphate ion transport (GO:0006817), indicating the common role of adaptive molecular evolution during alkaline adaptation in fish.

### Apoptosis and cell death

Extremely alkaline stress may cause extensive damage to fish, such as inducing cell apoptosis (Monteiro et al. 2009; Zhao et al. 2016, 2020). However, several fish species can survive in this harsh environment, such as schizothoracine (*Gymnocypris przewalskii*) (Tong, Fei, et al. 2017; Tong & Li 2020), Magadi tilapia (*Alcolapia grahami*) (Wilkie & Wood 1996), and Amur ide (*Leuciscus waleckii)* (Xu et al. 2017). This may also pose a barrier for alkaline tolerant fish as compared with alkaline intolerant fish. Significantly, positively selected *AIFM1, SH3RF1, YWHABA, PIK3R4* and *LAMP1* of alkaline tolerant species relative to alkaline intolerant species have enrichment in a set of apoptosis pathways, such regulation of apoptotic signaling pathway (GO:1902253) and macroautophagy (GO:0016236)(FIG. 5). Apoptosis is a form of programmed cell death that occurs in multicellular organisms, it plays a significant role in the biochemical events lead to characteristic cell changes (morphology) and death (Green 2011). Few direct evidence suggested the roles of these candidate genes in response to extreme alkaline stress in fish, but numerous studies had defined their functions in tolerance to harsh environments. *AIFM1* is a ubiquitous mitochondrial oxidoreductase involved in apoptosis, involved in sea bream (*Sparus aurata*) response to acute environmental stress (Bermejo-Nogales et al. 2014). *YWHABA* (14-3-3 protein beta/alpha-A) is an important gene showing the ability to bind a multitude of functionally diverse signaling proteins, such as transmembrane receptors (Fu et al. 2000), that involved in spotted sea bass (*Lateolabrax maculatus*) tolerance to saline stress (Zhang et al. 2019). In addition, *LAMP1* plays an important role in lysosome biogenesis and autophagy (Eskelinen 2006), which involved in common carp (*Cyprinus carpio*) response to hydrogen peroxide environment. Collectively, the presence of apoptosis related genes under positive selection may contribute to the alkaline adaptation of fish, how these adaptive molecular changes affect the ability programmed cell death remains unknown.

### Immune response

Extreme environments (including high pH) impact on the physiology of animals in a wide variety of ways. Recent advances in the understanding of environmental impacts were identified in relation to specific areas of immune function, such as increase in pH resulted in a general increase in immune function (Bowden 2008; Sridhar et al. 2020). Intriguingly, we identified different immune genes under positive selectin in alkaline tolerant species within highland (IL2RB, TLR8, IF144) and lowland (ILDR1 and CD44), but they all involved similar immune functions, such as inflammatory response (GO:0006954), natural killer cell activation (GO:0030101), and T cell mediated cytotoxicity (GO:0001913). In another word, we found different genes but conserved pathways may underlie fish adaptation to alkaline environment. IF144, IL2RB and ILDR1 are important components of toll-like receptor (TLR) signaling pathway that play key roles in the innate immune system (Rebl et al. 2010). For instance, our previous comparative studies in alkaline tolerant fish, *G. przewalskii* identified key genes involved in TLR pathway under selection (Tong, Fei, et al. 2017; Tong et al. 2015). In Nile tilapia (*Oreochromis niloticus*), another alkaline tolerant species and stands as ideal model to study extreme environment adaptation, a most recent comparative study identified numbers of immune genes involved in natural killer cell mediated cytotoxicity and NF-kappa B signaling pathway in response to alkalinity stress (up to pH = 8.9) (Zhao et al. 2020), and echoes our present results. Thus, it is possible that adaptive evolution changes (e.g. positive selection) acting on innate immune genes in alkaline tolerant species contribute to their adaptation to extremely alkaline environment.

### Energy metabolism

It is not surprising that we identified a set of genes under either accelerated evolution or positive selection, that enriched in diverse metabolisms, such as glucose metabolism, phosphate metabolism, sulfur amino acid metabolism and protein metabolism in alkaline tolerant species compared with their alkaline intolerant relatives. In general, metabolism processes were involved in fish response to varies environmental stresses (including alkaline stress) by a huge amount of research (Wood 1991). A most recent study highlights the genes involved in conserved mitochondrial pathways under selection in adaptation to extreme environment (Greenway et al. 2020). In addition to our previous studies in highland alkaline tolerant fish, we found a number of genes associated with energy metabolism processes under selection as well, such as mitochondrial function and protein metabolism (Tong, Fei, et al. 2017). Moreover, increasing studies on alkaline tolerant fish species (e.g. killifish) pointed out the roles of varied metabolisms in extreme environment (e.g. high pH) adaptation (Scott et al. 2019; Zhao et al. 2020; Wang & Guo 2019; Xu et al. 2017). Altogether, our analysis infer that the adaptive molecular evolution of metabolism associated genes may be indispensable and common features to alkaline adaptation as well as extreme environment adaptation in fish.

## Conclusion

In summary, this comparative genomics study of 15 fish species suggests the common role of alkaline adaptation in fish, regardless of highland or lowland background environments. Our results also highlight that the adaptive evolution of protein-coding genes are likely to play a crucial role in fish response to extreme environment, such as extremely high PH. Notably, this study provides putative genomic signatures of shift in selection and alkaline adaptation in several alkaline tolerant fish species, further study should include large scale of omics data of more alkaline tolerant fish species and their intolerant relatives as multiple pairs to demonstrate the genetic basis of alkaline adaptation in fish at genome-wide scale.

## Author Contributions

C.T. and K.Z. conceived this project. C.T. designed the project. C.T., Y.T. and K.Z. collected the samples. C.T. and M.L. performed the comparative genomics analyses. C.T. wrote the paper. All authors read and approved the final manuscript.

## Acknowledgments

We would like to thank four anonymous reviewers for helpful comments on this manuscript.

## Data Availability Statement

The Illumina sequencing reads have been deposited at NCBI Sequence Read Archive under the NCBI BioProject accession PRJNA684806.

## Supplementary table capture

supplementary table S1: Information for additional omics data.

supplementary table S2: Statistics of transcriptome assembly and protein-coding genes.

supplementary table S3: Statistics of orthologs in 15 fish species.

supplementary table S4: Patterns of molecular evolution in alkaline tolerant and alkaline intolerant species within lowland fish taxa and highland fish taxa.

supplementary table S5: Rapidly evolving gene repertoires in highland alkaline tolerant and lowland alkaline tolerant species.

supplementary table S6: Significantly enriched GO terms for rapidly evolving gene of highland alkaline tolerant and lowland alkaline tolerant species.

supplementary table S7: Positively selected gene repertoires in highland alkaline tolerant and lowland alkaline tolerant species.

supplementary table S8: Significantly enriched GO terms for positively selected gene of highland alkaline tolerant and lowland alkaline tolerant species.

supplementary table S9: Shared REG/PSG repertoires in highland alkaline tolerant and lowland alkaline tolerant species.

supplementary table S10: significantly enriched GO terms for shared REG/PSG of highland alkaline tolerant and lowland alkaline tolerant species.

